# A rod conformation of the *Pyrococcus furiosus* Rad50 coiled coil

**DOI:** 10.1101/2020.06.24.160820

**Authors:** Young-Min Soh, Jerome Basquin, Stephan Gruber

**Author notes:** Corresponding author, Stephan Gruber: S.G.

## Abstract

The Rad50-Mre11 nuclease complex plays a vital role in DNA repair in all domains of life. It recognizes and processes DNA double-strand breaks. Rad50 proteins fold into an extended structure with a ~20-60 nm long coiled coil connecting a globular ABC ATPase domain with a zinc hook dimerization domain. A published structure of an archaeal Rad50 zinc hook shows coiled coils pointing away from each other. Here we present the crystal structure of an alternate conformation displaying co-aligned coiled coils. Archaeal Rad50 may thus switch between rod-shaped and ring-like conformations as recently proposed for a bacterial homolog.

## Introduction

DNA damage occurs throughout the life of a cell. A particularly harmful type of DNA damage is the double-strand break (DSB). DSBs originate from intrinsic metabolic processes or from external sources. In some biological contexts, DSBs are also deliberately generated. If left unrepaired, DSBs may trigger cell death, while mistakes during DSB repair may lead to mutations and cancer.

One of the first repair factors localizing to DSBs is the Mre11-Rad50 nuclease complex, which processes broken DNA ends to initiate homologous recombination or nonhomologous end joining (Cejka, 2015; Kowalczykowski, 2015; Paull, 2018; Syed and Tainer, 2018). Rad50 proteins belong to the family of SMC-like proteins. They harbor a globular ABC ATPase domain, also called the head (Hopfner et al., 2000; Lammens et al., 2004). Rad50 proteins use a zinc hook—rather than a SMC hinge domain—to form homo-typic interactions with four cysteine residues, two from each monomer, coordinating a zinc ion (Hopfner et al., 2002; Park et al., 2017). The zinc hook and the head are distantly connected by an extended coiled coil ‘arm’ (Hohl et al., 2011). How the processing of DNA by the Mre11 (endo-)nuclease is restricted to broken DNA is poorly understood. A recent cryo-EM study on bacterial homologs however provided vital new insights (Kashammer et al., 2019).

The organization of Rad50 and SMC coiled coils has received significant attention in recent years. While the arms were initially regarded as flexible connectors, the emerging view is that they undergo coordinated structural transitions during the ATPase cycle for regulation and to perform work (Figure 1A). Two crystal structures of the Rad50 zinc hook report alternate coiled coil arrangement (Hopfner et al., 2002; Park et al., 2017). However, the two reported structures are of different origin limiting the possibility of direct comparison. In this study, we solved another crystal structure of the *Pyrococcus furiosus* (*Pfu*) Rad50 zinc hook. Unlike the previously solved structure with wide open coiled coils, the new structure revealed closely aligned coiled coils being more similar to the structure of the human Rad50 zinc hook and analogous to some structures of SMC protein fragments (Hopfner et al., 2002; Park et al., 2017; Soh et al., 2015). The new structure allows for a direct comparison of two states of a Rad50 arm at atomic resolution. Results from site-specific crosslinking performed in solution are consistent with the new rod model.

**Figure 1.**
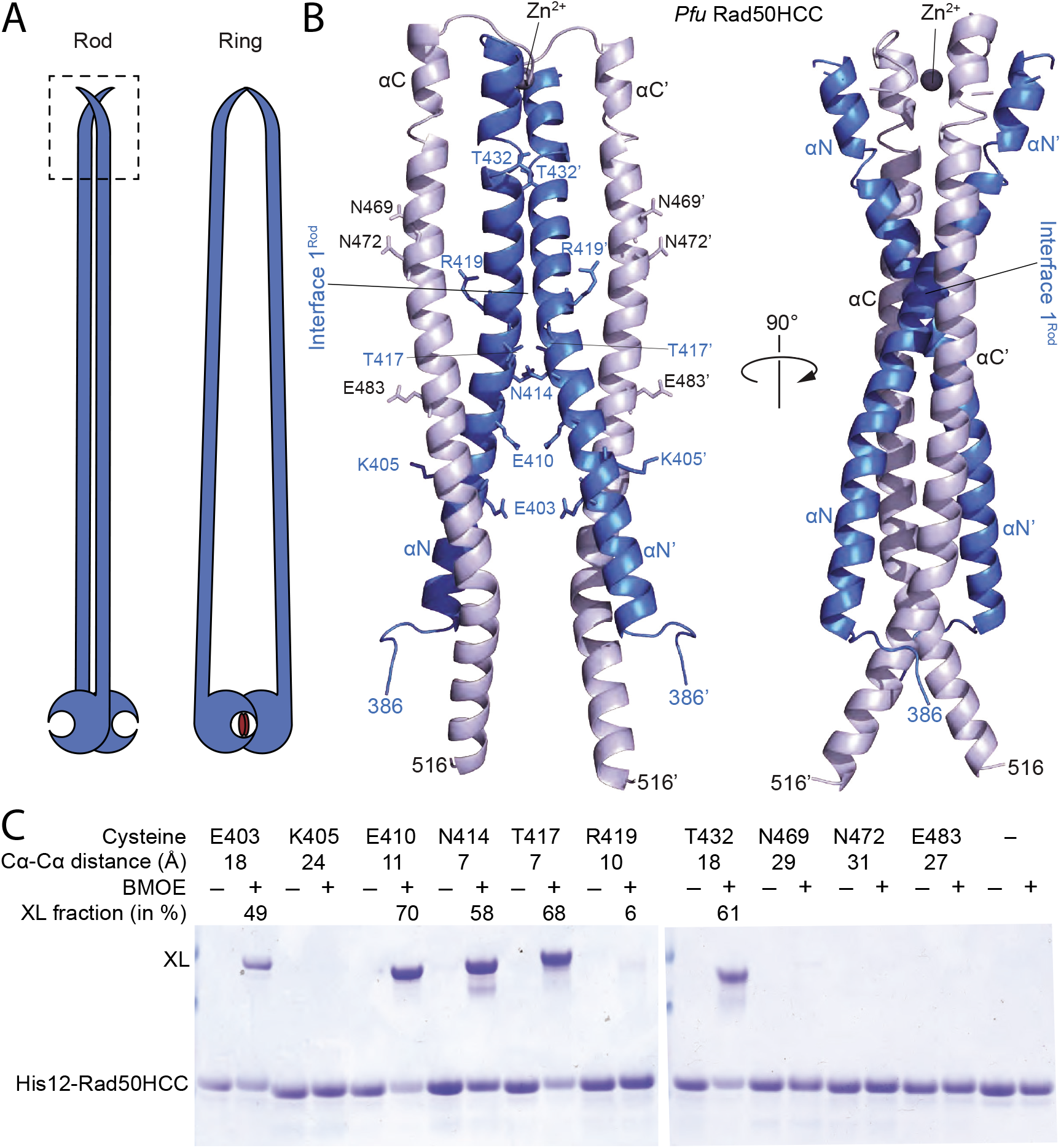
(A) Proposed organization of the Rad50 protein with disengaged heads (‘rod’, left panel) and with engaged heads (‘ring’, right panel). (B) Crystal structure of *Pfu* Rad50HCC showing two aligned monomers in front and side view in cartoon representation. Amino- (αN) and carboxy-terminal (αC) α-helices are displayed in blue and gray colors, respectively. Residues mutated to cysteine for cross-linking are denoted in stick representation. (C) BMOE cross-linking of His12-Rad50HCC variants. Cross-linked products were resolved by PAGE and stained by Coomassie. Cα-Cα distances of residue pairs are denoted together with estimated cross-linking efficiencies.

## Materials and Methods

### Protein Expression and Purification

For crystallization, the *Pfu* Rad50 zinc hook spanning residues 378-516 (‘Rad50HCC’) fused to a C-terminal cysteine protease domain (CPD) with a 10-residue histidine tag was expressed in BL21 DE3 Gold using ZYM-5052 auto-induction media at 24 °C for 24 hours. Cells were harvested by centrifugation, resuspended in lysis buffer (350 mM NaCl, 50 mM Tris/HCl pH 7.5, 3 mM β-mercaptoethanol), and sonicated. The supernatant was loaded onto a HisTrap column, which was then washed with five column volumes (CV) of buffer A (200 mM NaCl, 20 mM Tris/HCl pH 7.5, 3 mM β-mercaptoethanol). For elution, a linear gradient was applied using buffer A supplemented with 500 mM imidazole. Fractions corresponding to Rad50HCC-CPD-His10 were collected and treated with 1 mM phytate at 4 °C for 1 hour to cleave off the tag. The sample was diluted to 50 mM NaCl using buffer B (50 mM Tris/HCl pH 7.5, 3 mM β-mercaptoethanol) and then loaded onto a HiTrap Q column. For elution, a linear gradient was applied using buffer B containing 1 M NaCl. Fractions corresponding to Rad50HCC were collected and loaded onto a HisTrap column in order to remove the His-tagged CPD. The unbound fraction containing Rad50HCC was concentrated and loaded onto a Superdex 75 16/600 pg column, equilibrated in buffer C (100 mM NaCl, 20 mM Tris/HCl pH 7.5, 1 mM DTT). Fractions corresponding to the zinc hook were collected, concentrated to 25 mg/mL, flash frozen in liquid nitrogen, and stored at −80 °C.

The single cysteine mutants with an N-terminal 12-residue histidine tag were expressed and purified similar to the construct used for crystallization, with a few modifications. Instead of a HisTrap column, supernatant was loaded onto a gravity flow column containing HisPur cobalt resin. The column was washed with 5 CV buffer A followed by an additional 5 CV wash with buffer A supplemented with 5 mM imidazole. After eluting with buffer A, containing 150 mM imidazole, sample was loaded onto a HiTrap Q column and eluted as mentioned above. Fractions, corresponding to His12-Rad50HCC, were concentrated and loaded onto a Superdex 200 16/600 pg column, equilibrated with buffer C, containing 1 mM TCEP instead of 1 mM DTT. Protein was then concentrated, flash frozen in liquid nitrogen, and stored at −80 °C.

### Crystallization

The 25 mg/mL Rad50HCC sample was used to set up initial screens using the sitting-drop vapor-diffusion method at 4 °C and 25 °C. Initial crystals formed from precipitant solution containing 2.25 M ammonium sulfate and 50 mM Tris/HCl pH 8. These crystals were fished using nylon loops, soaked in mother liquor containing 30 % (v/v) ethylene glycol, and flash frozen in liquid nitrogen. Zn^2+^ signal was used to collect single-wavelength anomalous dispersion (SAD) data on the PXIII (06DA) beamline at the Swiss Light Source and processed using software in the XDS suite. The structure was deposited to the Protein Data Bank (PDB) under accession code: 6ZFF.

### *In vitro* BMOE cross linking

14.7 μM His12-Rad50HCC and single cysteine mutants were prepared in separate tubes containing 100 mM NaCl, 20 mM Tris/HCl pH 7.5, and 1 mM TCEP, on ice (Soh et al., 2019). BMOE crosslinker (20 mM stock in DMSO) was added to a final concentration of 1 mM. 10 min later, the reaction was quenched with β-mercaptoethanol (23 mM final). Loading dye was added to the samples, which were then incubated at 98 °C for 5 minutes and loaded onto Bis-Tris 4-12 % gradient gels. Bands were stained by CBB and relative band intensity was estimated using ImageJ.

## Results

### *P. furiosus* Rad50 zinc hook crystal structure revealing rod formation

We recently found that the *Pfu* zinc hook can functionally substitute for the SMC hinge domain in the bacterium *Bacillus subtilis* (Bürmann et al., 2017). Site-specific cross-linking of the chimeric SMC-Rad50 protein indicated that the zinc hook supports normal SMC coiled coil alignment. These results are difficult to reconcile with the open coiled coil organization observed in the published zinc hook structure (PDB: 1L8D) (Hopfner et al., 2002). We wondered whether the *Pfu* Rad50 zinc hook may readily adopt an alternate state with co-aligned coiled coils. To test this, we solved the crystal structure of the *Pfu* Rad50 zinc hook construct with adjacent coiled coil spanning residues 378 to 516, designated as Rad50HCC, at 3.0 Å. This construct harbors slightly longer coiled coils than the previously crystallized construct. Most of Rad50HCC was well resolved (residues 386 to 516), except few residues (441 to 446) in the central loop including one of the two cysteines that coordinate the zinc ion (Figure 1B). The unit cell contains a Rad50HCC dimer with the two monomers being positioned adjacent to one another in a way that co-aligns the coiled coils into a rod-like structure as previously observed for human Rad50 (PDB: 5GOX) (Park et al., 2017), but distinct from the open *Pfu* Rad50 zinc hook structure (PDB: 1L8D) (Hopfner et al., 2002).

### Site-specific cross-linking of zinc hook coiled coils

We used site-specific chemical cross-linking with the bis-maleimide compound BMOE to test whether *Pfu* Rad50 protein forms this rod-like organization also in solution. We engineered single cysteine mutants of a His12-tagged Rad50HCC variant based on the new structure. All mutant His12-Rad50HCC proteins behaved like the wild-type protein during expression and purification. Cross-linking levels were in excellent agreement with the distances between residue pairs in the dimer structure (Figure 1C). Most residue pairs that are closely located in the dimer (such as E410, N414, and T417) led to high cross-linking efficiency (70, 58, and 68 %, respectively) when mutated to cysteine, whereas residue pairs that are further apart in the structure (such as K405, N469, and N472) displayed hardly any cross-linking. R419C residues showed rather little cross-linking, despite the proximity in the dimer structure. The R419 sidechains are, however, facing away from one another. Hence the short BMOE spacer might not be able to bridge the two residues. The pattern of cross-linking is not explained by the arms-open structure (PDB: 1L8D) (Hopfner et al., 2002). In summary, the results obtained by chemical cross-linking correspond well with the new crystal structure, implying that it closely mimics a conformation observed in Rad50HCC and presumably also in native *Pfu* Rad50-Mre11 complexes.

## Discussion

The new structure of the *Pfu* Rad50 zinc hook shows that rod formation is not restricted to eukaryotic or human Rad50 proteins as previously suggested (Park et al., 2017) but widely conserved in Rad50 proteins (Figure 2A). At the zinc hook, four α-helices are arranged similarly in human and *Pfu* Rad50. The two amino-terminal α-helices cross each other at an open angle (^~^60 °) close to the zinc hook and form a dimer interface ‘1^Rod^’ (Figure 1B, 2A). A glycine residue is at the center of interface 1^Rod^ in both dimer structures. The surrounding residues have hydrophobic sidechains in the case of human Rad50 and small or charged sidechains in *Pfu* Rad50. The two carboxy-terminal helices build another interface (‘2^Rod^’) in the human Rad50 dimer (Figure 2A). The corresponding segments are, however, too distant to form a similar contact in *Pfu* Rad50. The *Pfu* Rad50 helices are instead involved in packing interactions, which possibly disrupt this interface in the crystal. In human Rad50, the carboxy-terminal α-helix features a kink at the location of interface 2^Rod^, while the amino-terminal segment at the same position includes a prominent non-helical disruption. Analogous deviations are absent from the *Pfu* Rad50HCC structure, albeit a few non-helical residues are present at the extreme amino terminus of the structure. While the structures may differ in some detail, their overall similarity provides further support to the notion that the rod conformation is not a crystal packing artefact and that it is conserved in distant Rad50 proteins (and in SMC proteins).

**Figure 2.**
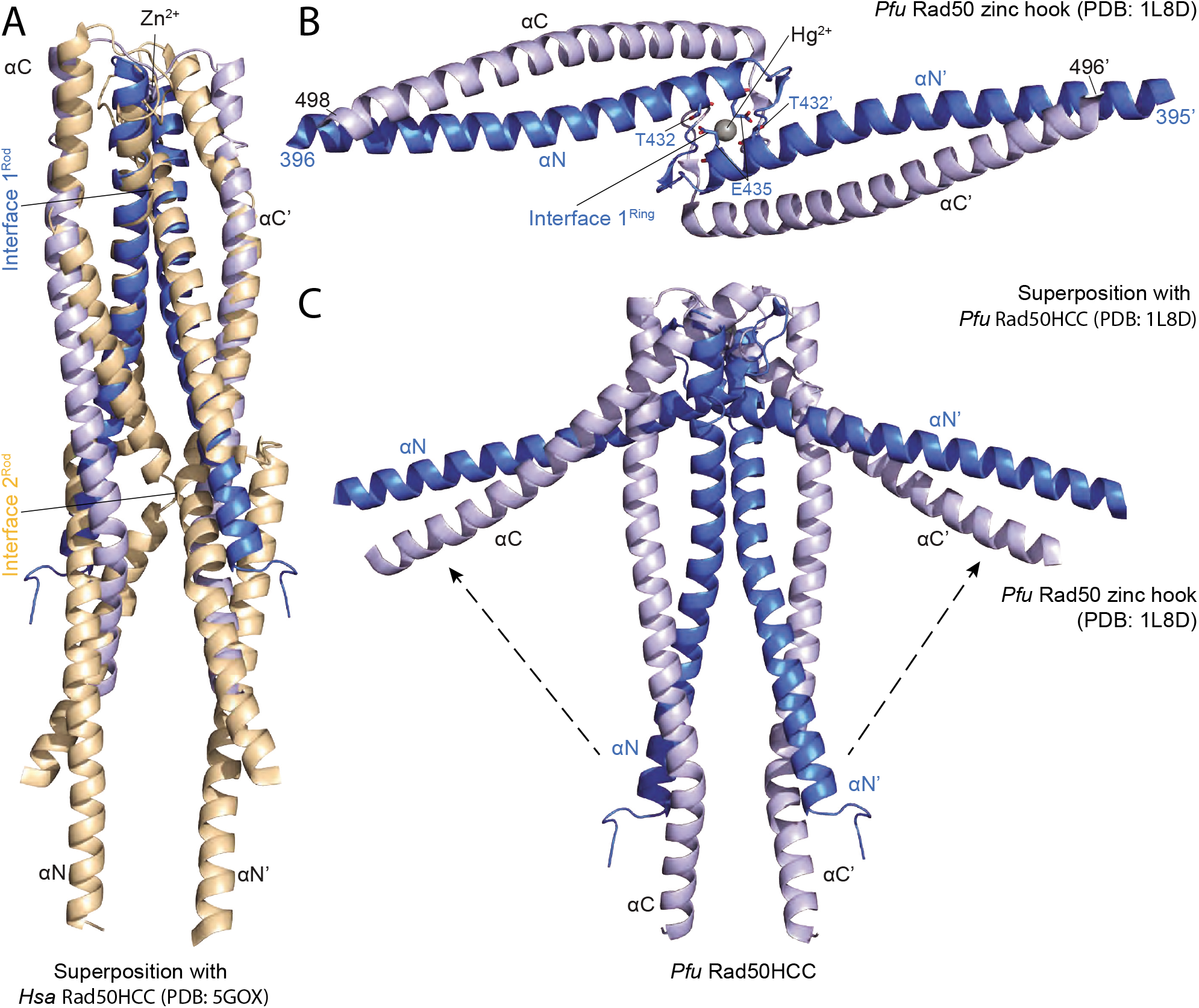
(A) Superimposition of *Pfu* Rad50HCC (in blue and gray colors as in Figure 1B) with human (*Hsa*) Rad50 zinc hook (in light orange colors). (B) Structure of *Pfu* Rad50 zinc hook in cartoon representation with residues forming interface 1^Ring^ highlighted in stick representation. (C) Superimposition of the two *Pfu* Rad50 zinc hook structures. Arrows indicate the tilting of the coiled coils.

Together with the previously published structure of the *Pfu* Rad50 zinc hook, the new structure implies that Rad50 coiled coils undergo a striking structural transition. Related opening and closure of SMC coiled coils has been proposed to be at the heart of the SMC motor for DNA loop extrusion (Diebold-Durand et al., 2017; Marko et al., 2019; Soh et al., 2015). A recent cryo-EM study on *E. coli* SbcCD, the bacterial homolog of Rad50-Mre11, observed a similar dynamic-yet-defined behavior of the Rad50 coiled coils and suggested that the transitions are key to DNA end sensing by Rad50-Mre11 (Kashammer et al., 2019). It identified two states of the Rad50 ATPase, one with open coiled-coil domains (‘scanning state’) and one with closed coiled coils (‘cutting state’). Only in the cutting state, the Mre11 nuclease is appropriately positioned for DNA processing and the cutting state can only form on broken DNA, together explaining end-specific DNA processing.

The coiled coils, however, were relatively poorly resolved in the cryo-EM maps of SbcCD (Kashammer et al., 2019). The atomic structures of the *Pfu* Rad50 zinc hook allow for a direct comparison of two states. The amino-terminal helix mediates inter-subunit contacts at the zinc hook also in the open form, albeit using an alternate interface (‘1^Ring^’), which is formed via hydrogen bonding (T432-E436’) and likely locks the α-helices in the wide-open arrangement (Figure 2B). Accordingly, the transition from the rod to the ring conformation involves disengagement of interface 1^Rod^ (and other putative rod interfaces), tilting of the coiled coils away from the rod axis by almost 90 ° each, and formation of interface 1^Ring^ (Figure 2C). Altogether, our data is consistent with the model for DNA end sensing by Rad50-Mre11 (Kashammer et al., 2019) and supports the notion that arm opening and closure is a fundamentally conserved feature of SMC and SMC-like proteins.

## Acknowledgements

We are grateful to the Crystallization Facility at the Max Planck Institute of Biochemistry and the Swiss Light Source (SLS). This work was supported by the Basic Science Research Program through the National Research Foundation of Korea (NRF) funded by the Ministry of Education (2017R1A6A3A03006597) and a Consolidator Grant (ERC COG 724482) by the European Research Council.

---

**Table 1.**
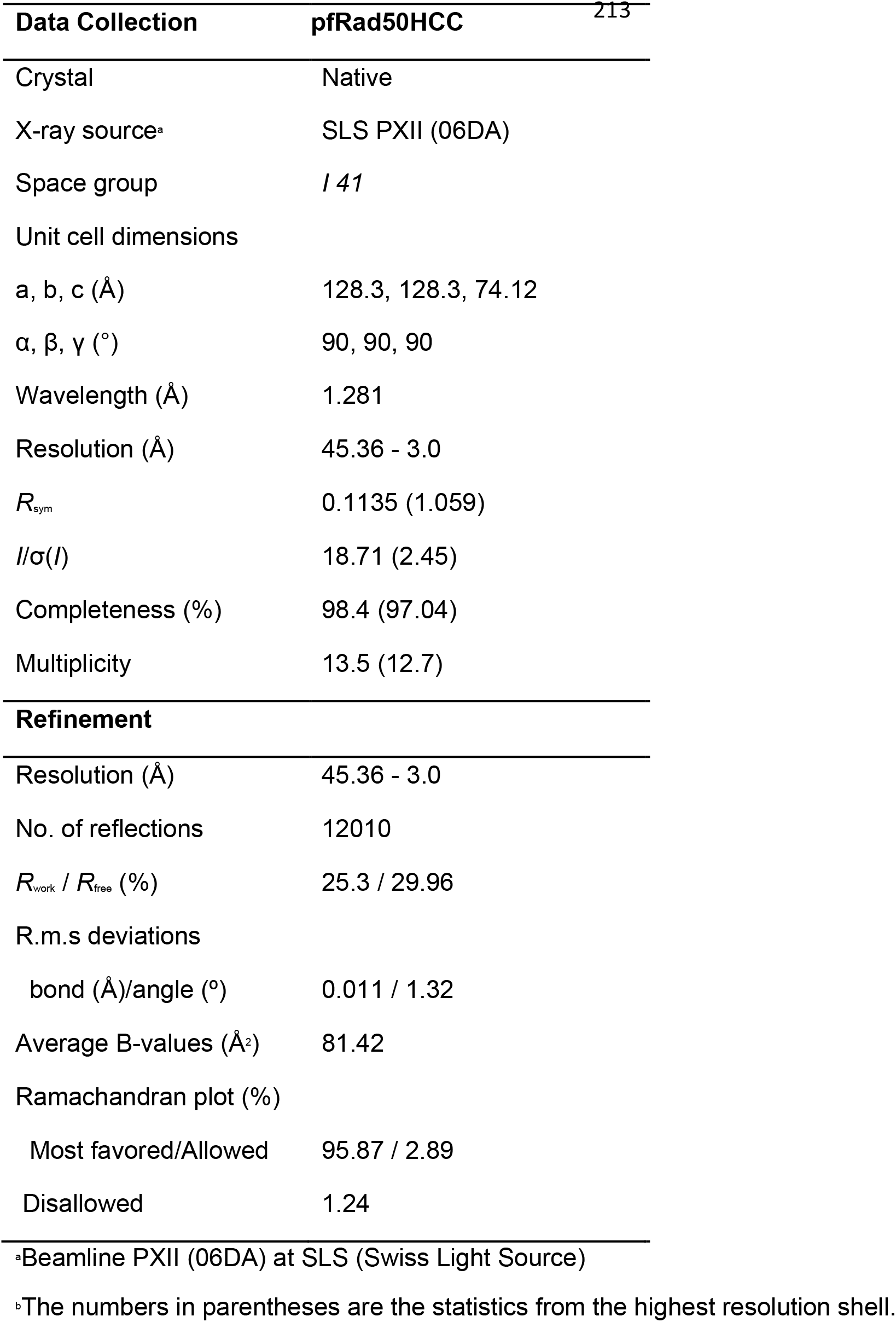
Data Collection and Structure Refinement Statistics

| Data Collection | pfRad50HCC | 213 |
| --- | --- | --- |
| Crystal | Native |  |
| X-ray source<sup>a</sup> | SLS PXII (06DA) |  |
| Space group | *I* 41 |  |
| Unit cell dimensions |  |  |
| a, b, c (Å) | 128.3, 128.3, 74.12 |  |
| α, β, γ (°) | 90, 90, 90 |  |
| Wavelength (Å) | 1.281 |  |
| Resolution (Å) | 45.36 - 3.0 |  |
| *R*<sub>sym</sub> | 0.1135 (1.059) |  |
| *I*/σ(*I*) | 18.71 (2.45) |  |
| Completeness (%) | 98.4 (97.04) |  |
| Multiplicity | 13.5 (12.7) |  |
| **Refinement** |  |  |
| Resolution (Å) | 45.36 - 3.0 |  |
| No. of reflections | 12010 |  |
| *R*<sub>work</sub> / *R*<sub>free</sub> (%) | 25.3 / 29.96 |  |
| R.m.s deviations |  |  |
| bond (Å)/angle (°) | 0.011 / 1.32 |  |
| Average B-values (Å<sup>2</sup>) | 81.42 |  |
| Ramachandran plot (%) |  |  |
| Most favored/Allowed | 95.87 / 2.89 |  |
| Disallowed | 1.24 |  |

<sup>a</sup>Beamline PXII (06DA) at SLS (Swiss Light Source)
<sup>b</sup>The numbers in parentheses are the statistics from the highest resolution shell.

## References

Bürmann, F., Basfeld, A., Vazquez Nunez, R., Diebold-Durand, M.L., Wilhelm, L., and Gruber, S. (2017). Tuned SMC Arms Drive Chromosomal Loading of Prokaryotic Condensin. Molecular cell.

Cejka, P. (2015). DNA End Resection: Nucleases Team Up with the Right Partners to Initiate Homologous Recombination. The Journal of biological chemistry 290, 22931–22938.

Diebold-Durand, M.L., Lee, H., Ruiz Avila, L.B., Noh, H., Shin, H.C., Im, H., Bock, F.P., Burmann, F., Durand, A., Basfeld, A., et al. (2017). Structure of Full-Length SMC and Rearrangements Required for Chromosome Organization. Molecular cell 67, 334–347 e335.

Hohl, M., Kwon, Y., Galvan, S.M., Xue, X., Tous, C., Aguilera, A., Sung, P., and Petrini, J.H. (2011). The Rad50 coiled-coil domain is indispensable for Mre11 complex functions. Nature structural & molecular biology 18, 1124–1131.

Hopfner, K.P., Craig, L., Moncalian, G., Zinkel, R.A., Usui, T., Owen, B.A., Karcher, A., Henderson, B., Bodmer, J.L., McMurray, C.T., et al. (2002). The Rad50 zinc-hook is a structure joining Mre11 complexes in DNA recombination and repair. Nature 418, 562–566.

Hopfner, K.P., Karcher, A., Shin, D.S., Craig, L., Arthur, L.M., Carney, J.P., and Tainer, J.A. (2000). Structural biology of Rad50 ATPase: ATP-driven conformational control in DNA double-strand break repair and the ABC-ATPase superfamily. Cell 101, 789–800.

Kashammer, L., Saathoff, J.H., Lammens, K., Gut, F., Bartho, J., Alt, A., Kessler, B., and Hopfner, K.P. (2019). Mechanism of DNA End Sensing and Processing by the Mre11-Rad50 Complex. Molecular cell 76, 382–394 e386.

Kowalczykowski, S.C. (2015). An Overview of the Molecular Mechanisms of Recombinational DNA Repair. Cold Spring Harbor perspectives in biology 7.

Lammens, A., Schele, A., and Hopfner, K.P. (2004). Structural biochemistry of ATP-driven dimerization and DNA-stimulated activation of SMC ATPases. Current biology : CB 14, 1778–1782.

Marko, J.F., De Los Rios, P., Barducci, A., and Gruber, S. (2019). DNA-segment-capture model for loop extrusion by structural maintenance of chromosome (SMC) protein complexes. Nucleic acids research 47, 6956–6972.

Park, Y.B., Hohl, M., Padjasek, M., Jeong, E., Jin, K.S., Krezel, A., Petrini, J.H., and Cho, Y. (2017). Eukaryotic Rad50 functions as a rod-shaped dimer. Nature structural & molecular biology.

Paull, T.T. (2018). 20 Years of Mre11 Biology: No End in Sight. Molecular cell 71, 419–427.

Soh, Y.M., Bürmann, F., Shin, H.C., Oda, T., Jin, K.S., Toseland, C.P., Kim, C., Lee, H., Kim, S.J., Kong, M.S., et al. (2015). Molecular basis for SMC rod formation and its dissolution upon DNA binding. Molecular cell 57, 290–303.

Soh, Y.M., Davidson, I.F., Zamuner, S., Basquin, J., Bock, F.P., Taschner, M., Veening, J.W., De Los Rios, P., Peters, J.M., and Gruber, S. (2019). Self-organization of parS centromeres by the ParB CTP hydrolase. Science 366, 1129–1133.

Syed, A., and Tainer, J.A. (2018). The MRE11-RAD50-NBS1 Complex Conducts the Orchestration of Damage Signaling and Outcomes to Stress in DNA Replication and Repair. Annual review of biochemistry 87, 263–294.

